# Simulations reveal unique roles for the FXR hinge in the FXR-RXR nuclear receptor heterodimer

**DOI:** 10.1101/2024.05.29.596427

**Authors:** Tracy Yu, Arumay Biswas, Namita Dube, C. Denise Okafor

**Author notes:** Authors contributed equally (Tracy Yu, Arumay Biswas).

## Abstract

Nuclear receptors are multidomain transcription factors whose full-length quaternary architecture is poorly described and understood. Most nuclear receptors bind DNA as heterodimers or homodimers, which could encompass a variety of arrangements of the individual domains. Only a handful of experimental structures currently exist describing these architectures. Given that domain interactions and protein-DNA interactions within transcriptional complexes are tightly linked to function, understanding the arrangement of nuclear receptor domains on DNA is of utmost importance. Here, we employ modeling and molecular dynamics (MD) simulations to describe the structure of the full-length farnesoid X receptor (FXR) and retinoid X receptor alpha (RXR) heterodimer bound to DNA. Combining over 100 microseconds of atomistic MD simulations with enhanced sampling simulations, we characterize the dynamic behavior of eight FXR-RXR-DNA complexes, showing that these complexes support a range of quaternary architectures. Our simulations reveal critical roles for the hinge in the DNA-bound dimer in facilitating interdomain allostery, mediating DNA binding and driving the dynamic flexibility of the complex. These roles have been hard to appreciate previously due to experimental limitations in studying the flexible hinge. These studies provide a much-needed framework that will enable the field to obtain a complete understanding of nuclear receptor quaternary architectures.

## Introduction

Nuclear receptors are a family of multidomain transcription factors whose gene regulation activity is modulated by lipophilic ligands^1^. The human genome contains 48 nuclear receptor genes^2^, broadly grouped into four classes by their mechanism of activation, including how they bind DNA. While Type I nuclear receptors are sequestered in the cytoplasm in the absence of a bound ligand^3,4^, Type II, III, and IV receptors reside in the nucleus regardless of ligand-binding status^1,5^. Type II receptors are the largest class, comprised of receptors that bind direct (DR), inverted (IR), and everted repeats (ER) of DNA as a heterodimer with retinoid X receptor alpha (RXR)^6,7^. A comprehensive understanding of the mechanisms underlying nuclear receptor function requires studying the structure and dynamic motions of receptors in functionally relevant contexts, e.g. in their dimeric and/or DNA-bound states.

Obtaining structural data on nuclear receptors in DNA-bound, dimeric states has been a long-standing challenge, largely due to the highly disordered regions within nuclear receptor domains that disrupt crystal packing^8–10^. Structures abound of the individual nuclear receptor domains, i.e the ligand binding domain (LBD) and DNA binding domain (DBD). These include dimeric LBD structures, portraying a dimerization interface between the two monomers, as well as DNA-bound DBD structures in both monomeric and dimeric states. In contrast, only three crystal structures have been reported depicting full-length Type II RXR heterodimers bound to DNA: PPARγ-RXR (DR1)^11^, LXRβ-RXR (DR4)^12^, and RARβ-RXR (DR1)^13^. These structures illustrate the complex architecture of full-length receptors and the difficulty of visualizing the intervening hinge linker between DBD and LBD. Indeed, the intrinsically disordered hinge is unresolved in five of the six nuclear receptor monomers represented in the three existing structures. Further, the three structures exhibit drastically different domain arrangements, emphasizing the need for dynamic characterizations of full-length receptors.

In our recent study, we utilized homology modeling and dynamics simulations for structural characterization of the full-length farnesoid X receptor (FXR)^14^. Our simulations predicted the existence of ligand-induced interdomain interactions, which we validated in experimental assays. Our past study revealed an important role for the hinge in mediating DBD-LBD communication. Importantly, we illustrated the dynamic nature of monomeric FXR, identifying a variety of structural arrangements adopted by the receptor. Here, we extend these approaches to model the dimeric, DNA-bound form of the FXR-RXR heterodimer. In this work, we vary both DNA and ligand identity, allowing us to sample conformations of the dimer in a range of functional states. We focus on features related to the overall architecture of the complex, building on the previously described structures of the dimerized individual domains (FXR-RXR LBD^15^ and FXR-RXR DBD^16^). We perform both long-scale (10-15 microseconds) trajectories and shorter (500 ns) classical and accelerated MD simulations, allowing us to probe multiple timescales and also achieve statistical rigor. Combined with functional assays, these studies uncover new roles for the FXR hinge in the context of the heterodimeric complex.

Our simulations of multiple DNA-bound FXR-RXR heterodimers reveal how the flexibility of FXR and RXR hinge regions allows the dimer to attain a range of conformational arrangements. The FXR hinge displays much higher flexibility than the RXR hinge. Regardless of their architectures, all structures conserve the dimerization interface between LBDs and DBD-DNA contacts reported in crystal structures^15,16^, confirming that the conformational dynamism of the full-length dimer does not compromise critical contacts needed for recognition. We do not observe any interactions between the FXR hinge and LBD as observed in our work on FXR monomer^14^ and as has been reported in other RXR heterodimers^11,13,17,18^, suggesting that interdomain allostery in FXR-RXR is distinct from monomeric FXR. However, we reveal an important role for the FXR hinge in interdomain communication, also observed in monomeric FXR, identifying a conserved feature between the two oligomeric states. Finally, simulations suggest a role for the FXR hinge in DNA binding. In summary, MD simulations are a powerful tool for describing the structure and dynamics of full-length, DNA-bound receptors, with the potential to yield valuable insights into transcription regulation mechanisms.

## Results

### The FXR-RXR dimer is conformationally dynamic

To characterize the structure of full-length FXR-RXR, we first generated a model of the heterodimer bound to the canonical response element, IR1. A DNA sequence containing the consensus IR1 sequence was used in the complex: 5’-CCG<u>AGGTCA</u>A<u>TGACCT</u>CG-3’ (repeated half-sites underlined). To model the DBDs bound to DNA, we used the crystal structure of the drosophila FXR-RXR homolog: EcR-RXR bound to IR1 (PDB 1R0N)^19^. We obtained the FXR-RXR LBD dimer from PDB 6A60^20^. To assemble both DBD and LBD dimers into a single dimeric complex, we used Modeller to generate the RXR and FXR hinge regions linking respective DBDs to LBDs **(Fig. 1A)**. In our default model, FXR is bound to the physiological ligand chenodeoxycholic acid (CDCA) while RXR is bound to its ligand 9-*cis* retinoic acid (9cRA).

**Figure 1.**
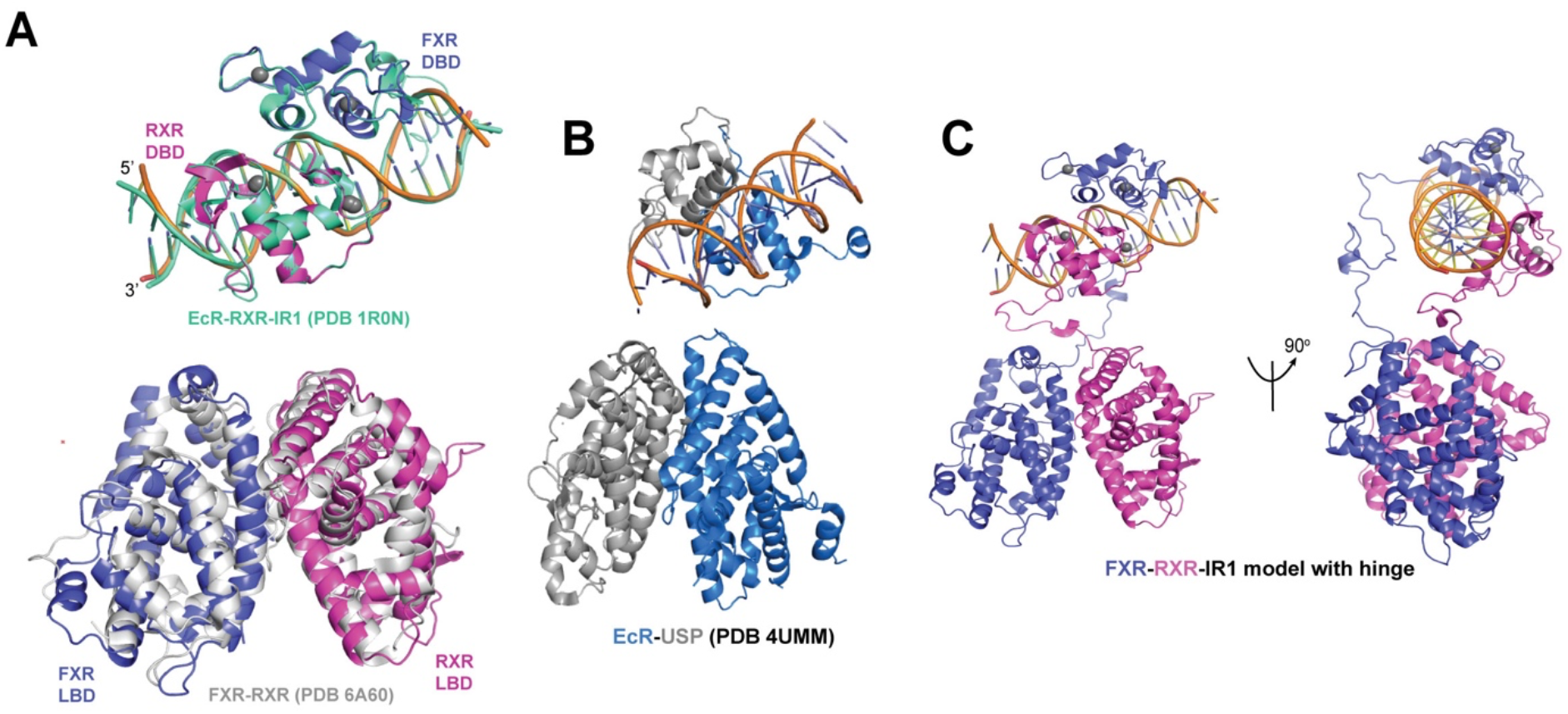
Structural model of FXR-RXR heterodimer. A) The FXR-RXR-DNA model was constructed by homology modeling. The DBDs bound to the DNA are based on the structure of the drosophila FXR-RXR homolog EcR-RXR (PDB 1R0N, green). The FXR-RXR LBD dimer is derived from PDB 6A60 (gray). Modeller was ultimately used to insert the hinge region between both FXR and RXR DBDs and LBDs. B) The full-length structure of the drosophila homolog EcR-USP complexed with IR1 (PDB 4UMM) provides the closest comparison for our structure, as the only IR1-bound full length experimental structure. C) Our FXR-RXR model allows visualization of the hinge, illustrating how a rotation can change the orientation of the LBDs relative to the DBDs.

Several interesting features emerge from our initial model. The RXR DBD binds on the 5’ end of the DNA, i.e. in the first half-site (5’-AGGTCA-3’) while FXR DBD binds in an inverted configuration at the 3’ end (5’-TGACCT-3’). The RXR DBD and LBD retain proximity to one another due to a compacted hinge. In contrast, the FXR hinge is hyperextended to connect between the FXR LBD and DBD at the 3’ end of IR1. Thus, this initial configuration excludes any contact between FXR domains and places the DNA close to the dimerized LBDs. This asymmetrical arrangement of full-length nuclear receptor dimers has been previously described^21^. Even though no full-length RXR heterodimers have been crystallized with IR1 DNA, the full-length structure of drosophila homolog EcR-USP is complexed with IR1, (PDB 4UMM)^21^ providing a reasonable comparison **(Fig. 1B)**. While the DBDs in both models share a similar arrangement, the LBDs are ‘inverted’ **(Fig. 1C)**, with the FXR and RXR hinges appearing to cross one another. Because our models allow visualization of the full hinge, which is not the case in crystal structures, we observe that small rotations in the FXR and RXR hinges would allow the LBDs in our model to readily shift into a similar state as observed in EcR-USP. The length and flexibility of both hinges would permit this rotation. Thus, even before running simulations, by merely visualizing the hinge in our models, we gain an appreciation for the dynamism of the DNA-bound dimer.

To gain a broad conformational overview of the FXR-RXR dimer, we constructed seven additional dimer models. To examine the effects of FXR ligands, we substitute obeticholic acid (OCA) and lithocholic acid (LCA) for CDCA to generate *OCA-IR1* and *LCA-IR1* respectively. To observe the effects of DNA sequence, we generate two models where the IR1 sequence in our original structure is replaced with the IR1 sequences taken from the promoter of FXR target gene phospholipid transfer protein (PLTP)^22,23^ and the ecdysone response element (ECRE)^23^, both of which are recognized by FXR (designated *CDCA-PLTP* and *CDCA-ECRE*). Finally, to compare apo- and ligand-bound states of the receptor, we toggled the ligand status of the heterodimer in three additional complexes: i) FXR-CDCA + apo-RXR (*apo-RXR)*, ii) apo-FXR + RXR-9cRA (*apo-FXR)*, iii) apo-FXR + apo-RXR (*apo-apo)*. For all eight models, we obtained MD trajectories ranging from 10-15 microseconds, enabled by the Anton 2 supercomputer^24^.

To assess the stability of the models, we performed RMSD calculations **(Fig. S1)**. Within the first microsecond, all complexes achieve a stable RMSD that persists until the end of the simulation. However, RMSD values reach anywhere between 2 and 20 Å before stabilizing, suggesting that the dimer complexes shift out of their initial conformation to adopt a range of stable states. To assess the structural integrity of the dimer complexes post-simulation, we asked how the LBD and DBD portions of the simulated models compare to crystal structures of FXR LBD and DBD dimers: PDB 6A60 and 8HBM **(Fig. 2A-E, Fig. S2)**. In nearly all cases, we observed excellent superposition of crystal structures with simulation structures. Alignment observed in the DBD dimers confirms that the zinc-finger motif is retained during simulation, as are all predicted contacts between the protein and DNA half-sites. Similarly, the LBD dimers retain their dimerization interface and orientation throughout the trajectories. We observe a few unaligned regions, which is a reasonable expectation for flexible regions that undergo dynamic perturbation during simulations. In summary, simulations reveal that while DNA-bound FXR-RXR dimer is conformationally dynamic, it retains expected dimeric contacts.

**Figure 2.**
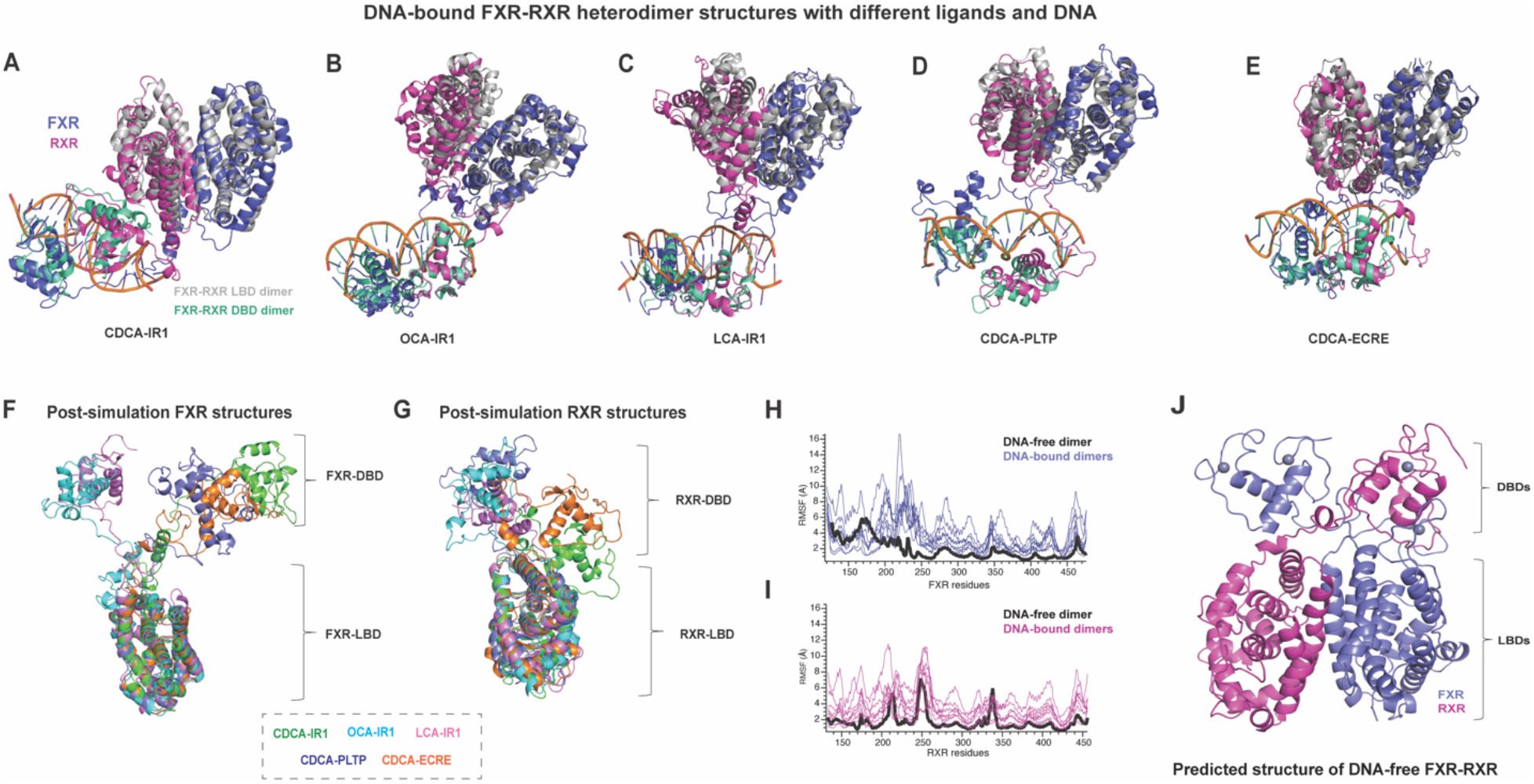
The FXR-RXR heterodimer is conformationally diverse. Post-simulation models of the DNA bound dimer are shown for A) CDCA-IR1, B) OCA-IR1, C) LCA-IR1, D) CDCA-PLTP, and E) CDCA-ECRE. A comparison between structures reveals that the dimer can achieve diverse conformational states. To assess how well the structures of the LBD and DBD dimers are preserved during simulation, each model is overlaid with the LBD and DBD crystal structures (shown in gray and green respectively). These comparisons confirm that individual domains retain their structure and preserve dimeric interactions during the simulation. F) Post-simulation models of FXR extracted from complexes reveal large conformational shifts in FXR. G) Post-simulation models of RXR extracted from complexes reveal smaller amplitude changes compared to FXR. H-I) A 15-microsecond MD trajectory was obtained for the DNA-free heterodimer complex, followed by an RMSF calculation. Comparison with DNA-bound dimer complexes reveals enhanced stability in both FXR and RXR in the absence of DNA. J) A predicted structure of the FXR-RXR heterodimer in the absence of DNA reveals that both FXR and RXR hinges collapse, leading to an overall compact and stabilized architecture.

### The FXR hinge mediates dynamic flexibility of the DNA-bound heterodimer

In our simulations on full-length monomeric FXR^14^, we observed that FXR adopts a wide range of interdomain architectures. To characterize the conformational variability of both FXR and RXR within the dimer, we extracted each monomer from all our post-simulation dimer structures and aligned the FXR and RXR monomers respectively by LBD. We observed a broad range of variability in FXR **(Fig. 2F)**, confirming that the DNA-bound dimer can undergo large motions. In comparison, RXR showed less variability, with a smaller distance between the LBD and DBD in all cases **(Fig. 2G)**. In summary, while the dimer can adopt diverse domain configurations, our models suggest that FXR undergoes large conformational adjustments to permit this dynamism, while RXR exhibits smaller changes.

The DNA elements that nuclear receptors bind to play important roles in shaping the architecture of full-length receptors. It has been demonstrated that IR versus DR DNA motifs influence both DBD and LBD arrangements in a full-length dimer^21^. To study how DNA influences the dynamics of the receptors more closely, we performed a 15-microsecond simulation of RXR-FXR in the absence of DNA. An RMSF comparison of DNA-bound versus DNA-free dimer reveals unexpected loss of stability in both FXR and RXR when DNA is present **(Fig. 2H-I)**. Further, our RMSD analysis of DNA-bound dimers **(Fig. S1)** showed that when partitioned by molecule (FXR vs RXR vs DNA), the DNA displays the smallest deviation in all complexes. Thus, in addition to being the most stable member of the complex, the DNA also stabilizes the protein. A close observation of the DNA-free dimer architecture reveals the origin of this effect **(Fig. 2J)**. In the absence of DNA, the FXR and RXR hinges adopt compact structures, bringing the DBDs and LBDs into greater proximity. These increased contacts between domains and hinges promote mutual stabilization and reduced movement. In contrast, the extended hinges of both FXR and RXR in the DNA-bound dimer allow a wider range of movement. While there is no experimental evidence for a DNA-free FXR-RXR heterodimer, this comparison reveals the indirect role of DNA in increasing the dynamism of the dimer, via extension of the hinge.

### The FXR hinge facilitates communication between FXR LBD and DBD

Our previous work in monomeric full-length FXR uncovered an important role for the hinge in mediating contact between the LBD and DBD^14^, as well as multiple DBD-LBD interfaces observed in simulations **(Fig. 3A)**. Crystal structures of full length RXR heterodimers have also shown physical contact between the two domains, e.g. in RARβ^13^. To quantify contacts between the LBD and DBD in the DNA-bound FXR-RXR heterodimer, we obtained Cα distance maps for our long-scale trajectories **(Fig. S3)**. Within a threshold of 20Å, no FXR LBD-DBD interdomain contacts are observed. To achieve sampling on longer timescales, we performed accelerated MD simulations on four complexes: CDCA-IR1, LCA-IR1, OCA-IR1, and apo-FXR. Triplicate 500-ns trajectories for each complex were combined and contacts were analyzed **(Fig. 3B)**. Even in these extended sampling conditions, we do not observe interactions between FXR LBD and DBD within the context of the dimer. This contrasts strongly with our observation in monomeric FXR.

**Figure 3.**
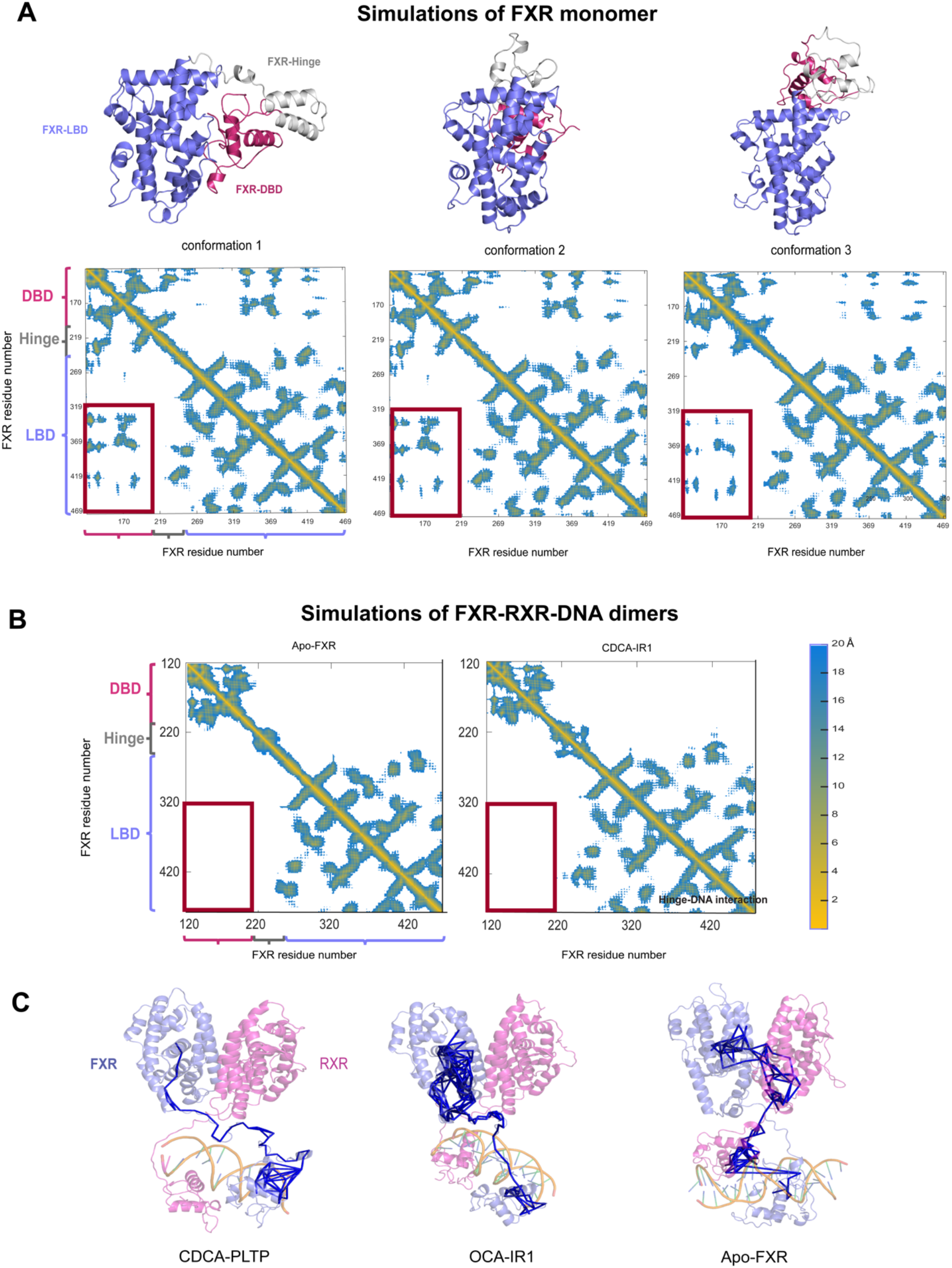
Interdomain FXR communication within the heterodimer. A) Three representations of monomeric FXR conformations previously reported^14^; interdomain contacts between LBD (in slate) and DBD (in magenta) are indicated within the red box, as shown in 9Cα distance matrices with a cutoff of 20Å. B) Cα distance matrices of FXR computed from accelerated MD simulations of the FXR-RXR dimer complex in the apo and CDCA-bound states. The matrices show that no LBD-DBD interdomain contacts present. Refer to **Fig. S4** for Cα maps of the full dimeric complexes. C) Suboptimal paths analysis for the FXR-RXR heterodimer complexes. Communication pathways from FXR LBD to DBD travels solely through the hinge when bound to FXR ligands but involves RXR residues in the apo state.

In the absence of physical contact between the two domains in our simulations, we turned to dynamic network analysis and suboptimal paths to identify amino acids mediating communication between the two domains. We obtained classical 500-ns MD trajectories in triplicate on holo- and apo-FXR DNA-bound dimer complexes, which were used for the suboptimal paths analysis. Strikingly, the FXR hinge is predicted as the mediator of interdomain communication, with 1583 and 197 paths traveling from LBD to DBD in OCA-IR1 and CDCA-PLTP complexes, respectively (See Methods). No involvement of RXR residues is observed. In contrast, suboptimal paths between FXR LBD and DBD in the apo-FXR complex incorporate residues from both RXR LBD and DBD. These results suggest ligand-specific trends within interdomain communication. In summary, while physical contact between FXR LBD and DBD is absent in the heterodimer, we still identify a role for the hinge in mediating communication between the two domains.

### Ligands drive conformational dynamics and function of the FXR hinge

The FXR hinge appears to play an important, ligand-specific role in mediating interdomain communication in the FXR-RXR heterodimer. This finding is consistent with our previous reports in the FXR monomer. To better understand the role of the FXR hinge, we quantified DNA-hinge interactions in our long-scale trajectories. We observe two unexpected interactions between DNA and hinge residues 197-244 **(Fig. 4A, B)**, with the closest contact occurring between residues 208 and 214. The hinge segment (208-KSKRLRK-214) maintains close proximity to the DNA backbone of base pairs 3-10, numbered as shown. We designate this region as the ‘KR patch’ due to the high frequency of lysine and arginine residues. These interactions are driven by electrostatic attraction between the positively charged residues in the KR patch and the negatively charged phosphate backbone. Unexpectedly, this hinge-DNA interaction is present in six of our eight FXR-RXR dimers, largely involving the same region of DNA (base pairs 3-9) **(Fig. 4B, S5)**. Notably, this region of DNA overlaps with base pairs 4-9 (AGGTCA), which is the DNA half-site that RXR binds. The RXR DBD binds in the major groove while the hinge KR patch occupies the minor groove.

**Figure 4.**
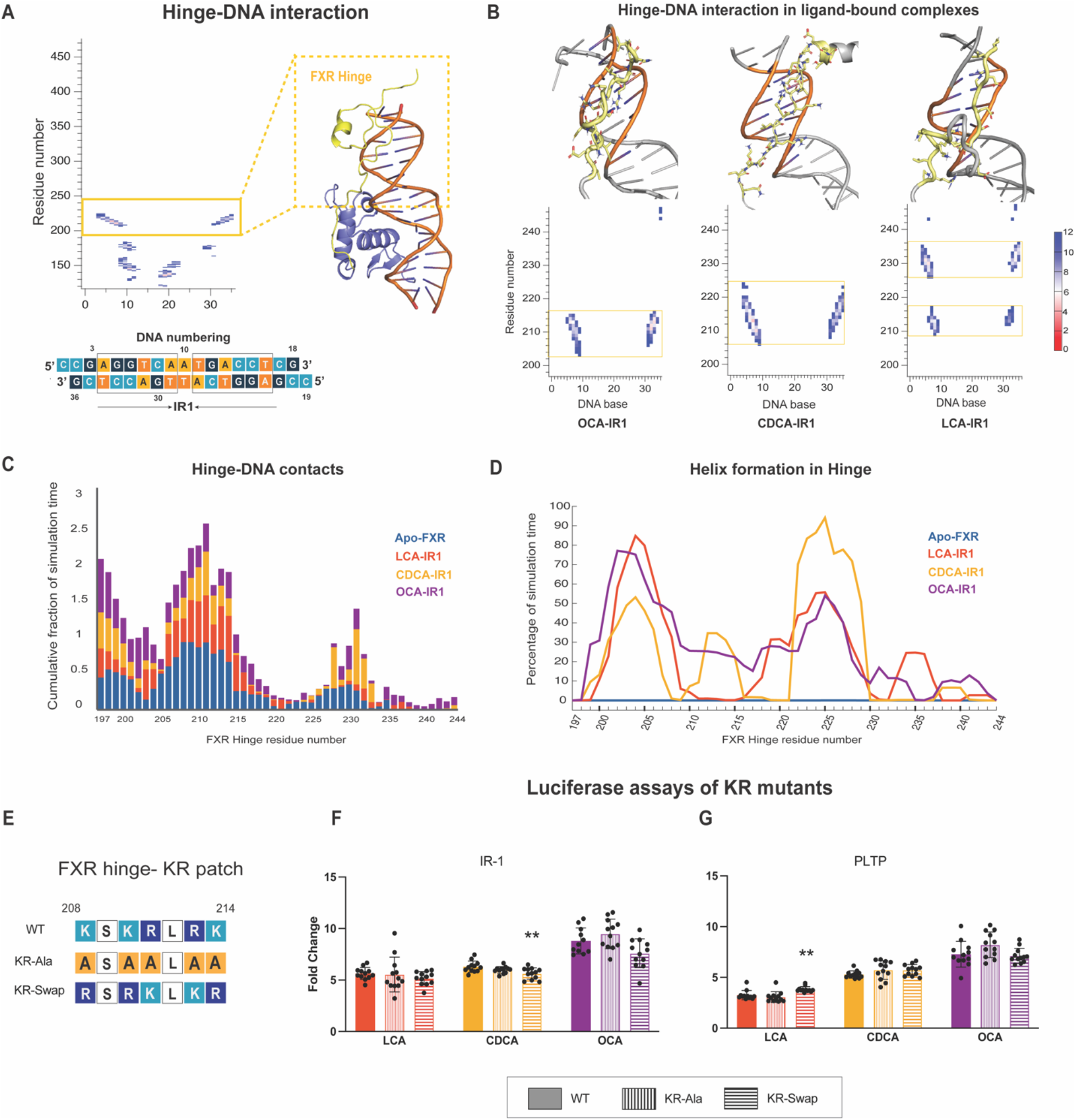
Interactions between FXR hinge and DNA. A) An FXR-DNA contact map is generated by identifying FXR Cα atoms and DNA C5’ atoms within 12 Å of each other. The contact map reveals interactions between the hinge and DNA, with IR1 DNA residues are labeled by numbering on contact map. B) FXR-DNA contact maps in three FXR-RXR dimer complexes are shown to illustrate hinge-DNA interactions. C) Fraction of simulation time during which the hinge residues contact DNA in various ligand-bound and apo FXR-RXR dimer complexes. D) Secondary structure analysis of the hinge. The data depicts the fraction of the simulation time during which each hinge residue is involved in helix formation. E) Basic amino acids in the KR patch were mutated to alanine (KR-Ala) or swapped (KR-Swap). F-G) Dual luciferase assays of the mutants were performed with response elements IR1 (F) and PLTP (G). Ligand concentrations used were 100 μM CDCA/LCA and 10 μM OCA. Wild-type FXR was used as the control for statistical significance, and results were analyzed using one-way ANOVA.

To explore the behavior of the hinge on enhanced time scales, we measured hinge-DNA interactions in our accelerated MD trajectories **(Fig. 4C)**. We observed consistent interactions between the KR patch region and DNA, a trend emerging in which the apo-FXR complex displays significantly higher protein-DNA interactions. This result suggests an influence of ligands on hinge dynamics, consistent with suboptimal paths analysis. Next, we quantified secondary structure in the FXR hinge of our aMD simulations **(Fig. 4D)**. Strikingly, the apo FXR dimer complex never attains a helical structure in its hinge, while significant amounts of helix formation are observed in holo-FXR dimeric structures. Thus, in addition to the observed ligand influence on the dynamics of FXR hinge, the data also suggest an inverse correlation between helical content of the hinge and its interaction with DNA, both modulated by ligands.

Based on our simulation results, we hypothesized that the KR patch of the FXR might play a role in activity. The nature of this role is unclear, as the presence of FXR ligands seem to reduce the involvement of the KR patch in DNA interactions. To determine whether there is physiological relevance of the KR patch, we created two mutants: KR-Ala and KR-Swap, and tested their activity in luciferase reporter assays **(Fig. 4E)**. The motivation behind mutating the basic amino acids to alanine (KR-Ala) was to disrupt the electrostatic interaction with DNA and observe its influence on transcription, the effect of which would be reversed by swapping lysines and arginines (KR-Swap). The assay was performed with two distinct FXR response elements, neither of which showed a significant reduction in the fold change compared to wild-type FXR **(Fig. 4F, G)**. Thus, in one respect, experiments support the notion that the non-specific interactions observed between the KR patch and DNA are not crucial for transcriptional activity. This result, however, does not eliminate the possibility of the KR patch’s role in DNA binding. Haelens et al reported a similar sequence (629-RKLKKLGN-636) in the androgen receptor hinge which, upon deletion, weakened binding with the androgen-selective response element (ARE) but increased transactivation by affecting other aspects of the transcriptional machinery^25^. Further experimentation will be required to validate the observed ligand-specific modulation of the structure and dynamics of the FXR hinge.

## Discussion

In this study, we sought to characterize the architecture and dynamics of the full-length DNA-bound FXR-RXR dimer. The application of in silico methods for studying full-length nuclear receptors is long overdue, as experimental approaches have been severely limited. To illustrate, while the gene for FXR was discovered in 1995^26^, it is only now, nearly 30 years later, that we have been able to visualize the structure of the full receptor, enabled by computational modeling^24^. While access to a supercomputer was required to obtain microsecond-scale simulations for complexes of this size, much longer (millisecond) timescales as well as replicate trajectories would be desirable to fully explore interdomain dynamics. Additionally, while we varied ligand and IR1 DNA sequences in our complexes, these timescales were not long enough to resolve unique effects induced by these substitutions.

Despite the challenges of applying classical MD simulations for studying large complexes such as those described here, we have gained valuable insight into the dynamic architecture of FXR-RXR. These DNA-bound complexes can adopt diverse conformations, enabled by the flexibility of the hinge regions, which can transition between extended and contracted forms **(Fig. 5)**. In spite of this conformational diversity, we do not see any interdomain contacts between FXR DBD and LBD, unlike what we observed in monomeric FXR. Nevertheless, the hinge is predicted to be the conduit of information between the two domains, a feature that appears to be conserved in monomeric FXR. The hinge is also predicted to mediate DNA interactions via the KR patch, a positively charged segment that interacts with DNA across multiple complexes. While experiments confirmed that this KR patch is not crucial for transcription, simulations suggest that ligands modulate the interaction between this hinge region and DNA. Thus, future efforts will be geared at elucidating the ligand-dependent function of the FXR hinge.

**Figure 5.**
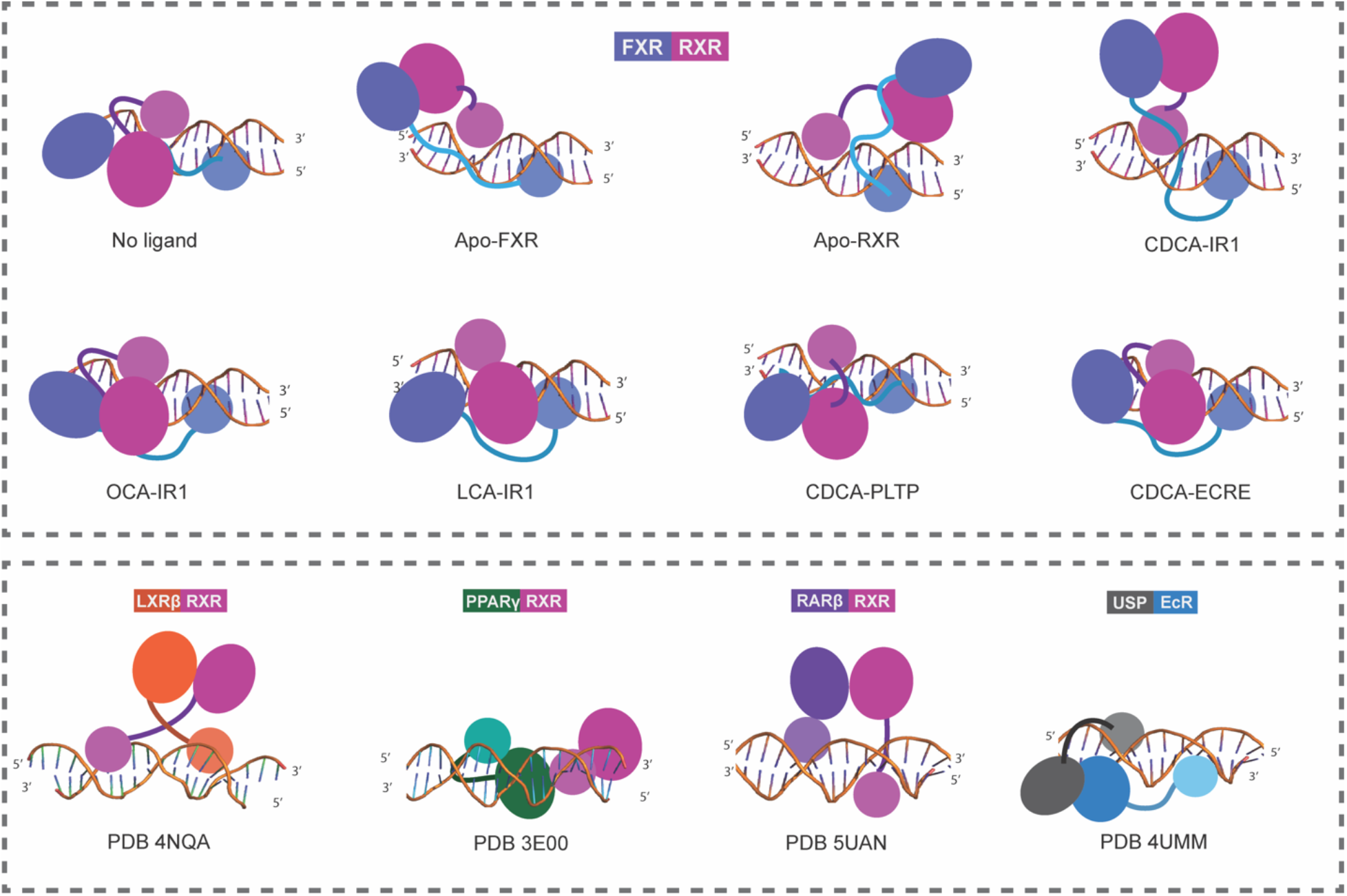
Comparison of quaternary architecture of FXR-RXR-DNA complexes and other RXR heterodimers. Schematic representations of post-simulation complexes from and existing RXR heterodimer structures (including drosophila homologs EcR-USP) reveal the diverse quaternary architectures nuclear receptors can adopt when bound to DNA.

Several questions remain regarding the architecture of nuclear receptor dimers, particularly from the perspective of understanding ligand-specific and DNA sequence-specific effects. Approaches such as cryoelectron microscopy, which can capture dynamic states of large complexes, will be valuable for achieving this insight. Regardless, simulation-based methods retain a powerful advantage by circumventing the challenges of sample preparation and handling, and other experimental logistics, which could become prohibitive for characterizing a complex in a large number of functional states.

## Materials and Methods

### Model development

To construct a model of full-length FXR-RXR heterodimer, we first used Modeller V9.23 to build a model of FXR DBD (residues R120-C196, Uniprot Q96RI1-1), which had not been crystallized at the time. This structure has since been solved^16^ and aligns with our model to RMSD < 2 Å. As a template, we used retinoic acid receptor alpha DBD taken from chain A of PDB 1DSZ^27^ (RARα:FXR DBD sequence similarity = 68%). We obtained a model of the RXR DBD (residues H133-Q206, Uniprot P19793-1) from PDB 4NQA^12^. We aligned both DBDs to the EcR-RXR dimer (PDB 1R0N)^19^ to generate the DNA-bound FXR-RXR DBD heterodimer. We obtained FXR-LBD (residues E245-Q476) and RXR-LBD (residues M230-T462) from PDB 6A60^20^. We separated both LBD and DNA-bound dimers in three-dimensional space by aligning them to the full-length LXRβ-RXR complex (4NQA)^12^. Modeller was then used to model the hinge linkers connecting each LBD to its corresponding DBD, the FXR-hinge (residues M197-K244) and RXR-hinge (residues E207-D229).

### Molecular dynamics simulations

Extended MD simulations were performed on Anton 2^28^. For binding energy calculations, triplicate 500 ns simulations were obtained using Amber18^29^ with GPU acceleration^30^. Antechamber^31^ from AmberTools27 was used to parameterize FXR and RXR ligands. The ff14SB forcefield^33^ and Generalized Amber Forcefield2^34^ were used for proteins and ligands, respectively. Complexes prepared for simulation on Anton 2 were solvated in a cubic box (103 × 98 ×77 Å^3^) of TIP3P water^35,^ with sodium and chloride ions added to reach a concentration of 150 mM NaCl. Complexes prepared for energy calculations were solvated in an octahedral box with a 10 Å buffer. All complexes were minimized, heated and equilibrated using the Amber18. Minimization was performed in four steps: i) with 500 kcal/mol. Å^2^ restraints on solute atoms, ii) 100 kcal/mol.Å^2^ restraints on solute atoms, iii) 100 kcal/mol. Å^2^ restraints on ligand atoms only, and iv) with no restraints on any atoms. Each minimization step utilized 5000 steps of steepest descent followed by 5000 steps of conjugate gradient. Heating to 300 K was performed using a 100-ps NVT simulation with 5 kcal/mol.Å^2^ restraints on all atoms. Pre-equilibration was performed in three 10-ns steps: i) with 10 kcal/mol.Å^2^ restraints on solute atoms, ii) with 1 kcal/mol.Å^2^ restraints on solute atoms, iii) with 1 kcal/mol.Å^2^ restraints on ligand atoms. After restraints were removed, Anton complexes were equilibrated for 50 ns before transferring to Anton 2 for extended MD. Complexes for energy calculations were simulated for 500 ns in triplicate. For all simulations, a 2-fs timestep was used with SHAKE. To evaluate the long-range electrostatics with particle mesh Ewald^30^ and Van-Der Waals forces, a 10-Å cutoff was used. CPPTRAJ^36^ was used to analyze RMSF, protein-DNA contacts, DSSP secondary structure analysis and Cα distance maps^37^. Network analysis and suboptimal paths were performed as previously described^38^.

For triplicate trajectories used in suboptimal paths analysis, the complexes above were simulated for 3x 500 ns. Source and sink for suboptimal paths were set to K148 in the FXR DBD and M332 on the FXR LBD respectively. Due to large numbers of paths between the two domains, the cutoff for suboptimal paths analysis was constrained to 2.

Accelerated MD trajectories were obtained following protocol we have previously described^14,39,40^. All accelerated MD trajectories were obtained in triplicate and combined prior to analysis.

### Mutant plasmids and dual luciferase assay

The amino acid residues 208-KSKRLRK-214 in the hinge region of FXR (Uniprot Q96RI1-1) were replaced with ASAALAA and RSRKLKR for KR-ala and KR-swap, respectively. Plasmids (vector pcDNA3.1+) expressing FXR mutants were obtained from Genscript.

HeLa cells were cultured in 10% fetal bovine serum (FBS)-supplemented Minimum Essential Medium alpha (MEMα) with 1% L-glutamine. 10000 cells were seeded in a clear, flat-bottom 96-well plate and allowed to grow for 24 hours before transfection. The cells were cotransfected with 5ng of FXR (wild type or mutant)-expressing plasmid, 5ng RXR-expressing plasmid, 50ng of minimal promoter (minP)-driven firefly luciferase reporter plasmid, and 1ng of SV40-driven renilla luciferase reporter plasmid (internal control) using Fugene HD (Promega). The response element IR1 and PLTP were in the minP promoter upstream of the firefly luciferase gene. 24 hours after transfection, the cells were treated with RXR ligand 9cRA and FXR ligands LCA, CDCA or OCA. The final concentration of the ligands inside the wells was 10μM for 9cRA and OCA, and 100μM for LCA and CDCA. After incubation for 24 hours, the firefly and renilla luciferase activities were quantified using the Dual-Glo® Luciferase Assay System (Promega) and Spectramax iD5 plate reader. Transactivation was calculated as fold change in luciferase activity with respect to DMSO, which was the vehicular control.

All experiments were performed in three biological replicates, with four technical replicates for each. To analyze statistical significance, the dataset was subjected to a one-way ANOVA test for comparing the mutants against the wild-type for each ligand in GraphPad Prism V10.

## Acknowledgments

Anton 2 computer time was provided by the Pittsburgh Supercomputing Center (PSC) through Grant R01GM116961 from the National Institutes of Health. The Anton 2 machine at PSC was generously made available by D.E. Shaw Research. C.D.O. is funded by an NSF award (CAREER: 2144679).

