## Supporting Information for "Simulations reveal unique roles for the FXR hinge in the FXR-RXR nuclear receptor heterodimer"

*Authors contributed equally

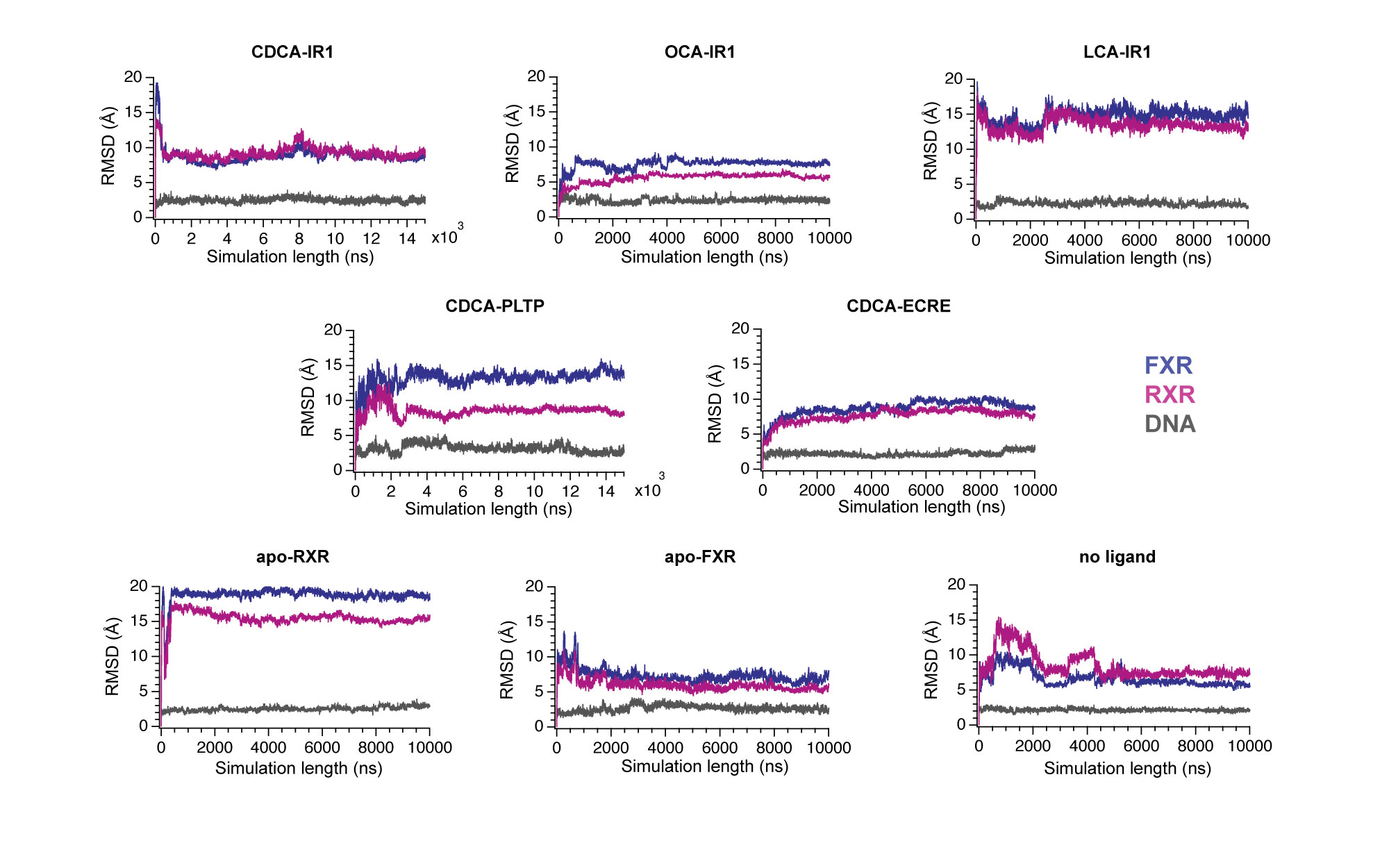

 Figure S1. RMSD analysis reveals that all complexes undergo an initial conformational change during simulation before stabilizing.

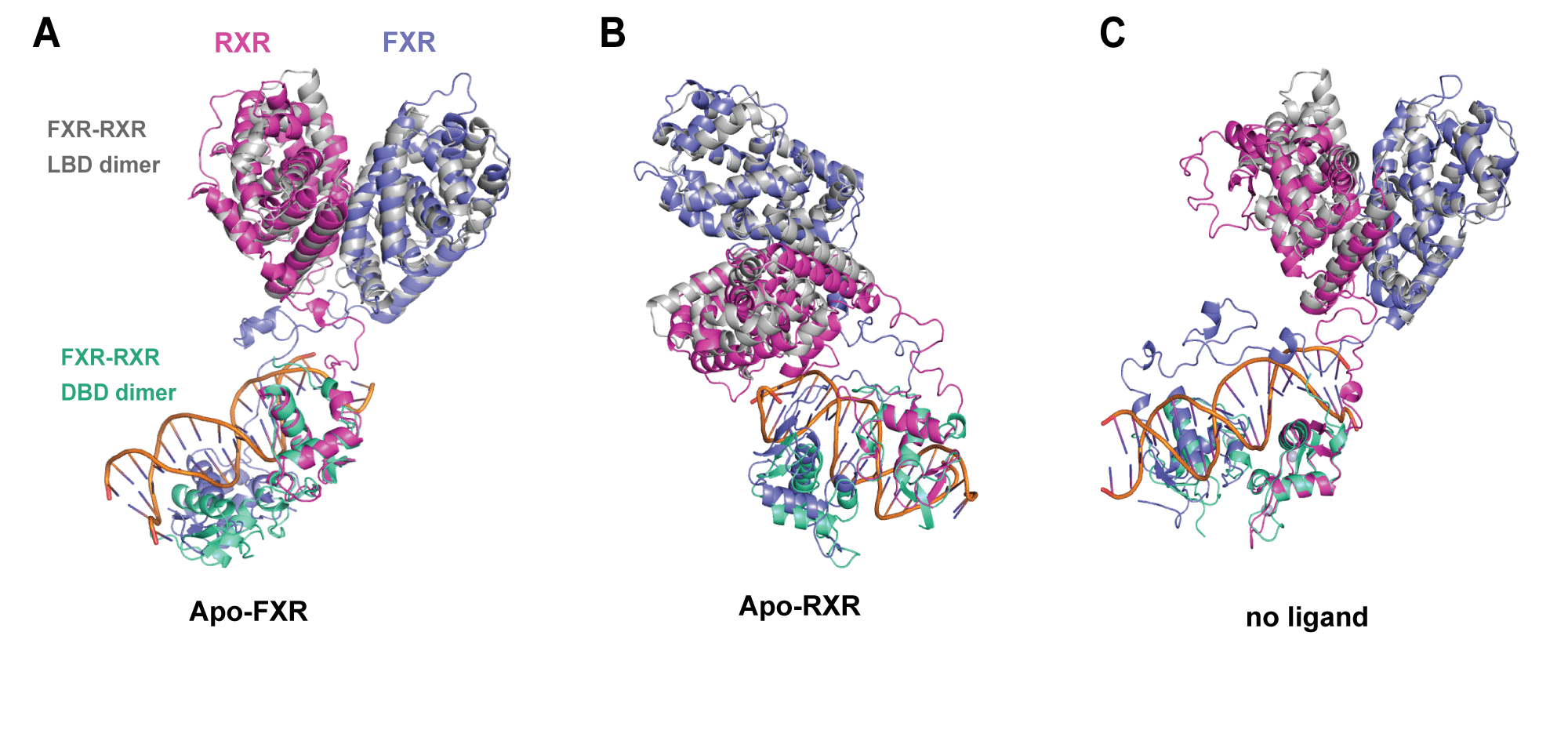

Figure S2. Post-simulation models of the DNA bound dimer are shown for A) apo-FXR,  B) apo-RXR, and C) no ligand complexes. Each model is overlaid with the LBD and DBD crystal structures (shown in gray and green, respectively). These comparisons confirm that individual domains retain their structure and preserve dimeric interactions during the simulation.

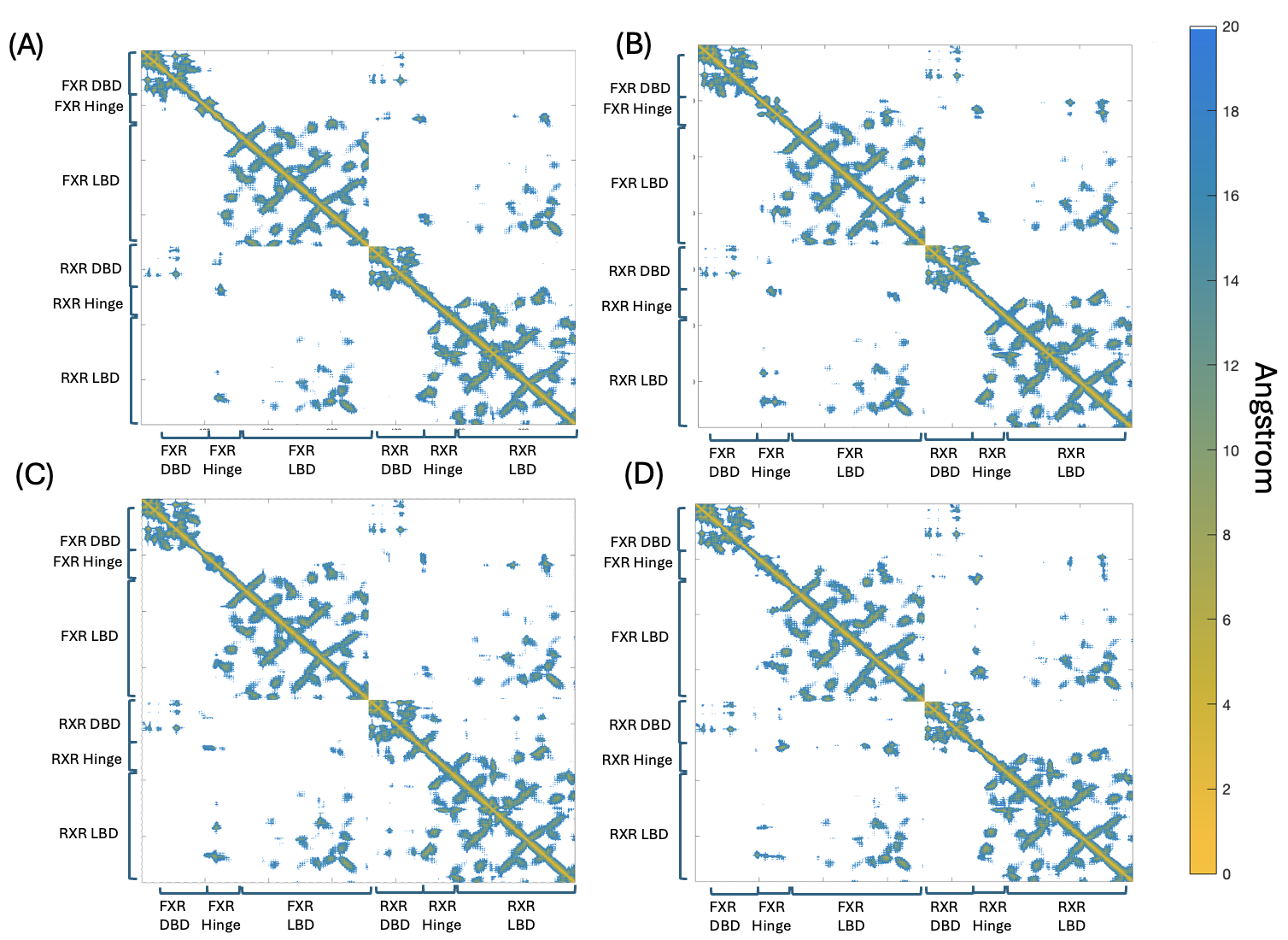

Figure S3. Cα distance maps of the full dimer complex A) apo-FXR,  B) LCA-IR1, C) CDCA-IR1, D) OCA-IR1 as analyzed from the 15μs long scale MD simulations. We do not observe any FXR LBD-DBD interactions here.

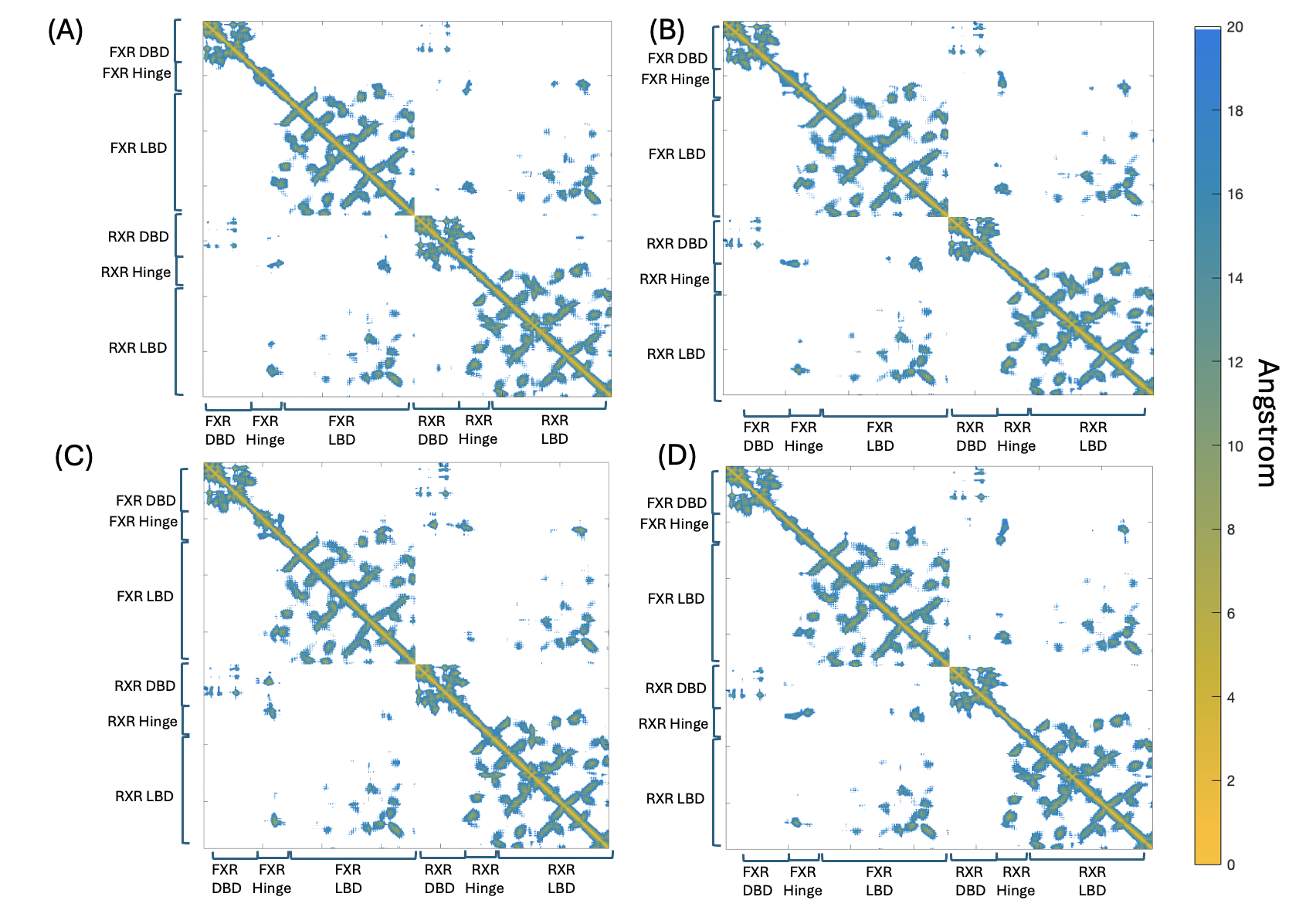

Figure S4: Cα distance maps of the full dimer complexes A) apo-FXR,  B) LCA-IR1, C) CDCA-IR1, D) OCA-IR1 as analyzed from the accelerated MD simulations. We do not observe any FXR LBD-DBD interactions here.

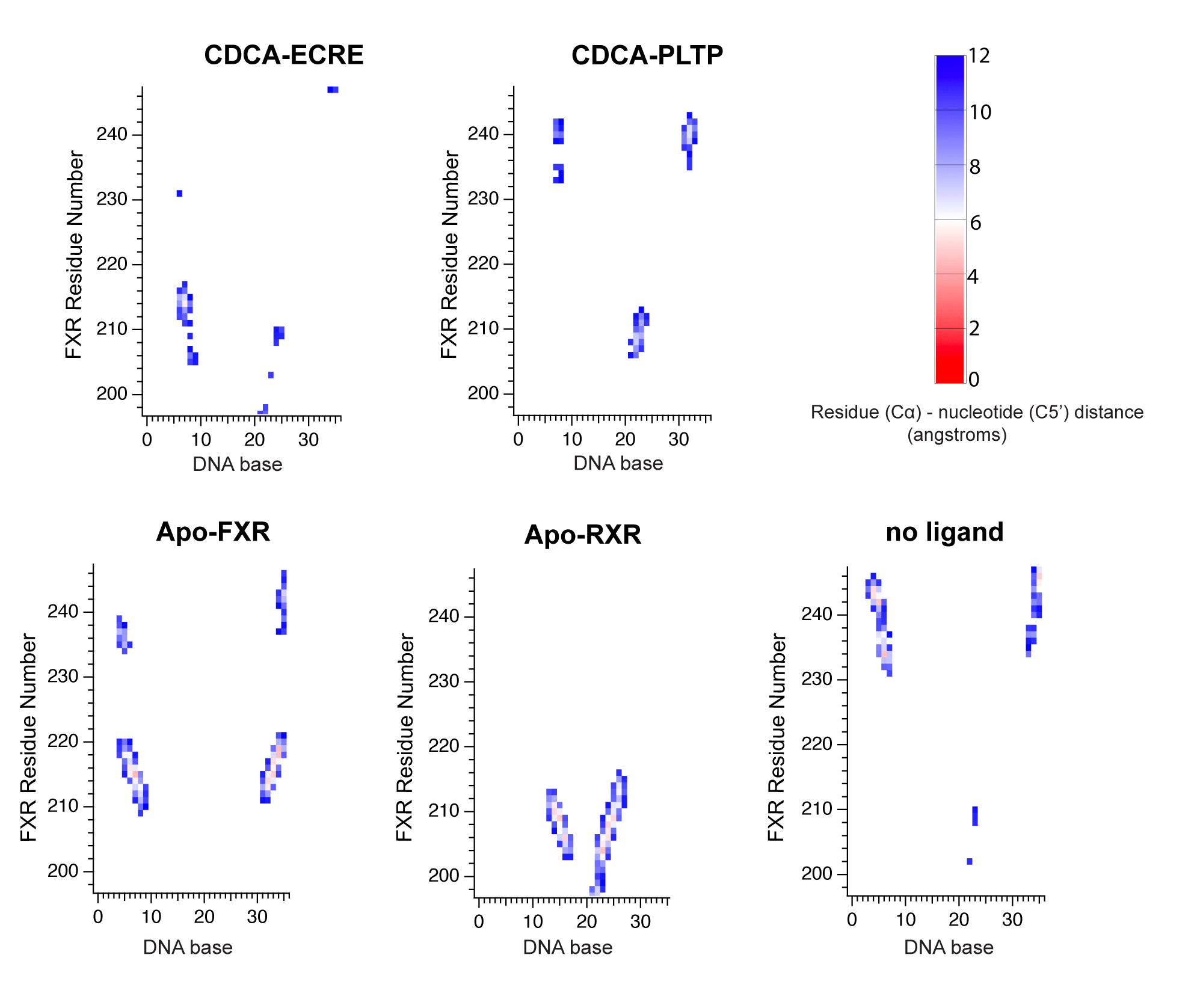

Figure S5. FXR-DNA contact maps are generated for complexes by identifying FXR hinge Cα atoms and DNA C5’ atoms within 12 Å of each other. The contact maps reveal hinge interactions with the KR patch (residues 208-214) as well as other regions of the hinge.
